# Chlorpyrifos induces lung metastases and modulation of cancer stem cell markers in triple negative breast cancer model

**DOI:** 10.1101/2024.11.04.621565

**Authors:** Marianela Lasagna, Mariana Mardirosian, Daniel Zappia, Lucia Enriquez, Noelia Miret, Lara Dahir, Elsa Zotta, Andrea Randi, Mariel Núñez, Claudia Cocca

## Abstract

Breast cancer is a major public health problem, and distant metastases are the main cause of morbidity and mortality. Chlorpyrifos is an organophosphate that promotes Epithelial-Mesenchymal Transition-like phenotype in breast cancer cell lines and modulates the Breast Cancer Stem Cells activating two key processes related to the metastatic cascade. Here, we investigated whether Chlorpyrifos may induce distant metastases in an *in vivo* triple negative tumor model. Also, we studied the expression of Breast Cancer Stem Cell and Epithelial-Mesenchymal Transition activation-markers in Triple Negative Breast Cancer mice tumors and human cells. We demonstrate that Chlorpyrifos modulates stem cell plasticity as a function of growth conditions in monolayer or three-dimensional culture. Furthermore, Chlorpyrifos decreased the doubling period, increased tumor volume, stimulated the infiltration of adjacent muscle fibers and induced lung metastases in mice. Finally, Chlorpyrifos modulated the expression of Epithelial-Mesenchymal Transition and BCSC markers in mice exposed to the pesticide. All our findings confirm that Chlorpyrifos promotes breast cancer progression, enhances stemness and Epithelial-Mesenchymal Transition marker expression and generates lung metastases in an *in vivo* model induced in mice.

## 1. Introduction

Breast cancer is a major public health problem, affecting more than 1.6 million women worldwide. Its incidence is on the rise in most countries and is expected to increase further in the next 20 years, despite current efforts to prevent this disease (Sung et al., 2021). In Argentina, breast cancer was the most commonly diagnosed malignant tumor in both sexes in 2020 and accounted for 32% of all cancer diagnosed in women (IARC, 2020). The incidence rate of breast cancer was the highest among all types of malignant tumors, with 73 cases per 100,000 population, as was the mortality rate, with 18.9 cases per 100,000 population (IARC, 2020).

The prognosis among patients with breast tumors is usually excellent when the cancer is diagnosed in its early stages, with a 5-year survival rate of 99%. Unfortunately, breast cancer cells can migrate to distant sites along the body, especially to the lung, liver, bone, and brain, in a process known as metastases, which is the leading cause of death. Less than 20% of breast cancer patients with distant metastases survive after five years (Fares et al., 2020; Jin et al., 2019). Metastatic disease is the primary cause of cancer morbidity and mortality and is responsible for about 90% of cancer deaths (Chaffer and Weinberg, 2011). Although many types of cancers are initially susceptible to chemotherapy, over time they can acquire resistance through diverse mechanisms, such as DNA mutations and metabolic changes that promote drug inhibition and degradation (Housman et al., 2014).

Exposure to environmental chemicals is ubiquitous, and there is growing evidence proposing that these factors may increase the risk of breast cancer development and progression (Kolak et al., 2017). Chlorpyrifos (CPF) is an organophosphate pesticide that has been widely used in our country and in the whole world until 2023. Oral, inhalation, and dermal exposure are the major contributing exposure pathways for this pesticide. First, CPF is one of the most widely detected pesticides in fruit and vegetables, so it is the main source of non-occupational exposure, generating important consequences for human and wildlife health (McLachlan, 2016; Thomas Zoeller et al., 2012). Second, inhalation constitutes another relevant exposure route for agro-applicants, rural workers, and family members with whom they live in rural areas.

We previously demonstrated that CPF causes serious endocrine-related adverse effects in mammary gland (Ventura et al., 2016, 2012), and increases tumor incidence with a minor latency period in N-nitroso-N-methylurea-induced mammary tumors in rats (Ventura et al., 2019). Recently, we demonstrated that CPF is able to promote epithelial-mesenchymal transition-like phenotype in MCF-7 and MDA-MB-231 breast cancer cell lines and modulates the breast cancer stem cells (BCSC) pool activating two key processes related to the initiation of the metastatic cascade (Lasagna et al., 2022, 2020).

Epithelial-mesenchymal transition (EMT) is described as a cellular program characterized by a group of phenotypic changes including loss of epithelial cell profile, enhanced cell mobility, and acquisition of aggressiveness. Extended literature shows that environmental and synthetic chemicals such as benzo(a)pyrene, a polycyclic aromatic hydrocarbon that can act as an AhR agonist (Yoshino et al., 2007), nicotine (Yu and Chang, 2013), cadmium and chromium (Chakraborty et al., 2010; Ding et al., 2013), pesticides (Zucchini-Pascal et al., 2012) or other endocrine-disrupting compounds (Palacios-Arreola et al., 2022) have the potential to induce cancer metastases through regulation of EMT markers and activation of migratory processes. These hazardous chemicals are able to regulate many transcription factors such as Slug and several signaling pathways mediated by Transforming Tumor Growth Factors β, Wnt/β-catenin, NOTCH, HEDGEHOG, NF-kappa-B pathways and tyrosine kinase receptors (Lee et al., 2017).

Aforementioned pathways are crucial for cancer stem cells self-renewal and maintenance (Takebe et al., 2011). These cells constitute a small percentage (0.05–1%) of tumor cells defined by the expression of stem cell marker genes such as OCT4 (Octamer-Binding Transcription Factor 4), SOX2 [(Sex Determining Region Y)-box 2)], Nanog, and ALDH1A1 (Aldehyde dehydrogenase family 1 member A1). BCSC were first described by Al Hajj et al in 2003 and were characterized by expressing the adhesion molecule cluster of differentiation (CD) 44, but not CD24. It was previously published that the injection of a few numbers of CD44^+^/CD24^-^ cells was enough to induce a rapid induction of tumors in mice (Al-Hajj et al., 2003). CD44 is a transmembrane glycoprotein whose major ligand is hyaluronic acid and additionally, it may act as a co-receptor mediating HER family and MET receptor tyrosine kinase signaling (Yang et al., 2011). Although CD44 was initially proposed to be a simple stemness marker, there is growing evidence that this protein promotes breast cancer progression by modulating signaling cascades and enhancing interactions between CD44 and the extracellular matrix (Zhao et al., 2016). CD24, a ligand of P-Selectin, is a small cell surface protein like mucin. While CD24 low expression has been proposed as a marker of BCSC, many studies show that its overexpression in various human cancers is an important marker of malignancy and bad prognosis (Altevogt et al., 2021; Huth et al., 2021). Furthermore, it is suggested that CD24 is expressed at a higher level in progenitor cells or metabolically active cells and lower in highly differentiated cells (Fang et al., 2010). Bakal et al. reported a novel role of CD24 at the interface of the immune system and tumor cells. CD24 expression may determine the response to immunotherapies because elevated expression may inhibit macrophage function (Barkal et al., 2019).

It is now clear that BCSC are involved in tumor development, cell proliferation, metastatic dissemination and resistance to chemotherapy and radiotherapy (Batlle and Clevers, 2017; Chang, 2016; Peitzsch et al., 2017). Hu et al. have shown that prostaspheres that are enriched in prostate cancer stem cells and express estrogen receptors alpha (ERα) and beta (ERβ) as well as GPR30, proliferate in response to low doses of estradiol and the endocrine disruptor Bisphenol A (Hu et al., 2012).

Finally, angiogenesis is another process intimately related to the development of distant metastases. It is a process that involves the formation of new vessels from preexisting ones, which are critical for solid tumor growth and invasion (Park et al., 2014). Vascular endothelial growth factor (VEGF-A) is a predominant activator of endothelial cell functions and one of the most important proangiogenic factors secreted by tumor cells (Chi et al., 2015). It was demonstrated that Hexachlorobenzene increases angiogenesis and VEGF-A expression in a xenograft model with MDA-MB-231 (Pontillo et al., 2013) and MCF-7 (Zárate et al., 2020) human breast cancer cells. CD31 or Platelet and Endothelial Cell Adhesion Molecule 1 (PECAM 1) is highly expressed by vascular endothelial cells and is involved in angiogenesis, immune escape, and tumor growth (Abraham et al., 2018). CD34 is commonly used as a marker for hematopoietic stem cells and hematopoietic progenitor cells (Sidney et al., 2014).

In short, all the above allows us to hypothesize that the environmental pollutant CPF may promote breast cancer progression by amplifying the BCSC pool. To prove our hypothesis, we designed our experiments to investigate whether CPF is able to induce distant metastases in an *in vivo* triple negative tumor model. To understand the mechanism involved in CPF transforming-actions, we studied the expression of BCSC and EMT activation-markers in primary tumors. We also evaluated the effect of CPF exposure on ALDH1A1, CD24 and CD44 stemness parameters and Vimentin, β-catenin, and Slug EMT markers in TNBC human cells using both bi– and tridimensional cultures.

## 2. Material and Methods

### 2.1. Cell culture

The estrogen-independent cell line MDA-MB-231 is used as a model of TNBC, poorly differentiated and highly malignant. They were purchased from the American Type Culture Collection (ATCC, USA). Cells were maintained as we previously reported (Ventura et al., 2015). The CPF concentrations used for all experiments were 0.05 and 50 μM. For the experiments, CPF was diluted in 0.5% ethanol (Vehicle-Merk, USA).

MDA-MB-231 cells were grown in monolayer and exposed to CPF (0.05 and 50 μM) for 72h. After treatment, mammosphere formation assay was performed using the protocol described in our previous publication (Lasagna et al., 2022).

### 2.2. Animals and treatment

Six-to eight-week-old female N: NIH (S) athymic mice (La Plata Laboratory Animal Facility, Buenos Aires, Argentina) were maintained at 22-23 °C room temperature, humidity close to 56% and light-dark cycles of 12 h of duration. Female N: NIH (S) athymic mice were housed into germ free environmental conditions. Animals were maintained for a week to allow its acclimatization in the new room. Animals were kept following both the guidelines for Care and Work with Laboratory Animals (U.S. National Academy of Sciences, 2011) and the ARRIVE guidelines (Animal Research: Reporting of *In Vivo* Experiments); the corresponding ARRIVE Compliance Questionnaire is enclosed as complementary material. The protocol was approved by the Institutional Committee for the Care and Use of Laboratory Animals of the Faculty of Pharmacy and Biochemistry, UBA (Exp-UBA N°0062405/15 and renewed Exp-UBA N° 20557/19).

CPF was dissolved in corn oil and then administered orally three times a week. Control animals only received corn oil. The doses used for the experiments were selected according to the international limits in force at the time of initiating this research. For the assessment of risk, we use the doses associated with toxicological exposures at the moment the experiments were performed: 0.001 and 0.1 mg/Kg/day, corresponding to the Acceptable Daily Intake (ADI) and No Observed Adverse Effect Level (NOAEL), respectively (European Food Safety Authority, 2019).

#### 2.2.1 MDA-MB-231 xenograft model in N: NIH (S) athymic female mice

For tumor induction, 7×10^6^ MDA-MB-231 cells were inoculated subcutaneously (s.c.) into the right flank of each mouse from a batch of 3 randomly selected animals. When the tumors reached a volume of 500 mm^3^, the mice were euthanized, and the tumors were excised. A randomly selected tumor fragment of approximately 4 mm^3^ was implanted subcutaneously under sterile conditions in each of the mice anesthetized with a mixture of ketamine/xylazine intraperitoneally (80-100 mg/kg + 5-10 mg/kg, respectively). Once xenografts reached a diameter of 0.5 cm, the mice were randomly separated into three different groups (Control; CPF 0.001 and CPF 0.1; *n=3*) to start the intoxication.

#### 2.2.2. Tumor growth and histopathology analysis

To evaluate tumor growth, the animals were checked for a lapse of 22 days, three times a week and two perpendicular diameters were measured with a caliper. Tumor size was calculated as we previously described (Ventura et al., 2019). After euthanasia, the primary tumors were measured, removed, and weighed. Liver, regional lymph nodes and lungs were also removed for metastasis studies. The samples were collected and processed as we described (Cocca et al., 1998). Collagen-Masson’s trichrome staining (TRI) was performed following the protocol described in Jones (2020).

#### 2.2.3. Immunohistochemical assay

PCNA (Proliferating Cell Nuclear Antigen), β-catenin, Slug and CD44 protein expression and localization were determined by immunohistochemical technique as we previously described (Ventura et al., 2016). Briefly, after deparaffinization, antigen retrieval was performed. Then, the slides were incubated overnight at 4 °C with primary antibodies using mouse anti-PCNA (1:100, Santa Cruz Biotechnology, USA), mouse anti-β-catenin (1:50, Santa Cruz Biotechnology, USA), rabbit anti-Slug (1:100, Invitrogen, USA) and mouse anti-CD44 (1:100, Santa Cruz Biotechnology, USA). Immunoreactivity was detected by using horseradish peroxidase-conjugated anti-polyvalent IgG (Vectastain Universal Elite ABC Kit, Vector Laboratories, USA) according to manufacturer specifications, and visualized by 3,3-diaminobenzidine staining (Sigma Chemical Co., MO, USA). Finally, the specimens were counterstained with hematoxylin and microscopy observation was performed with an optical microscope (Nikon Eclipse E200, Japan).

#### 2.2.4. Quantitative real-time PCR analysis

MDA-MB-231 cell line grown in monolayer or as mammospheres as well as a fragment of the primary tumor were homogenized in Quick-Zol (Kalium Technologies, Argentina) for 5 minutes, for RNA isolation and to allow complete dissociation of the proteins as we previously described (Miret, 2016). Total RNA was quantified by measuring optical densities at 260 and 280 nm. cDNA was synthesized from the RNA template via reverse transcription using random hexamer primers and MMLV Reverse Transcriptase (Promega Corporation, USA) according to the manufacturer’s instructions. Real-time PCR (qRT-PCR) was used to analyse the relative expression levels of CD44, CD24, ALDH1A1, Slug, Vimentin, CTNNB1 and β-actin. Primers are listed in ***Table 1***. The specificity of each primer set was monitored by analysing the dissociation curve. Relative mRNA quantification was performed by ΔΔCt method using β-actin as the housekeeping gene.

**Table 1.**
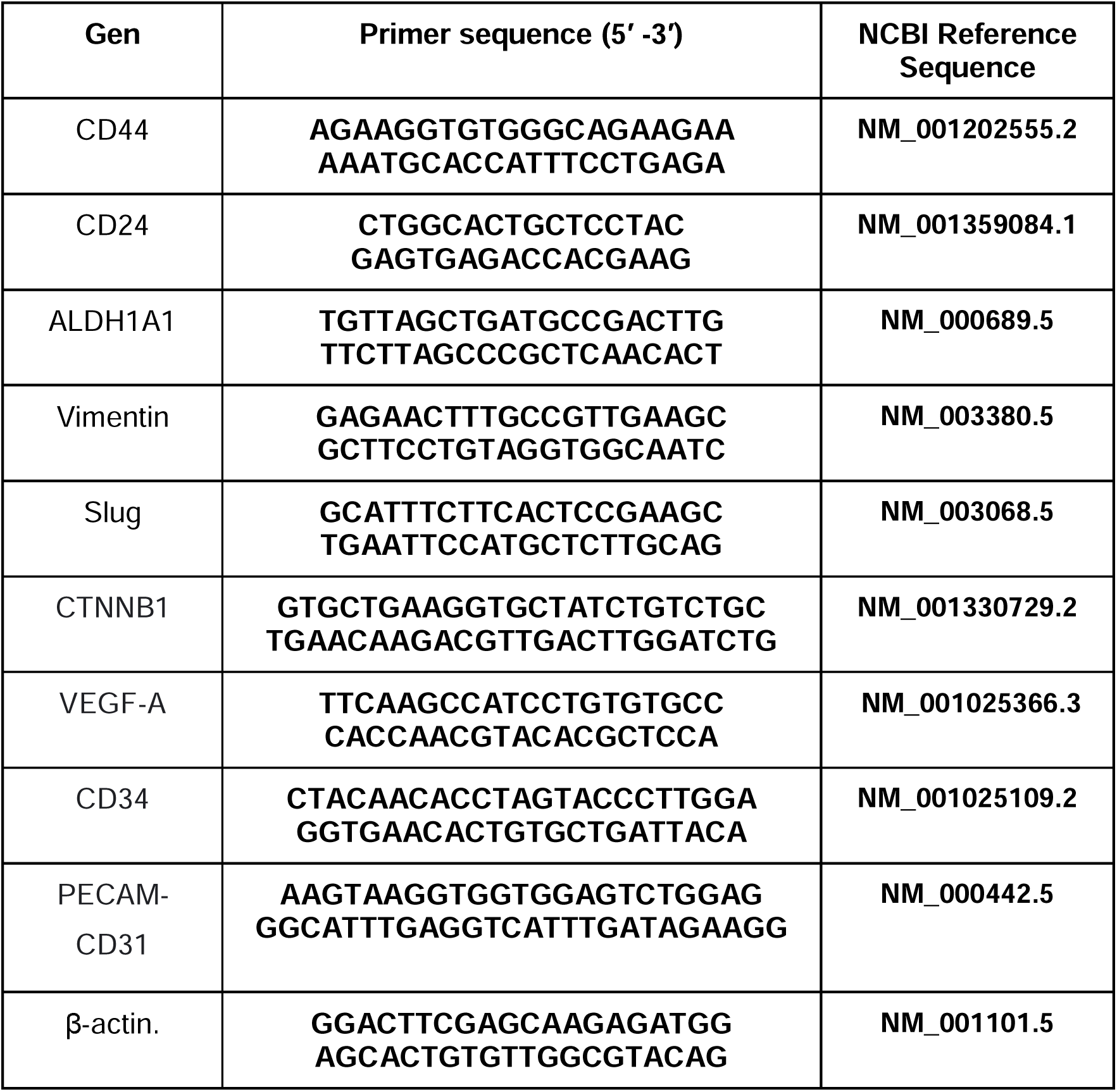
List of Primers Sequences used for RT-qPCR.

## 3. Statistical Analysis

Graph Pad Prism 8.0.1 (GraphPad Software Inc., USA) was used for data analysis. The statistical test performed is indicated in the legend of each figure. *p* values under 0.05 were considered statistically significant.

## 4. Results

### 4.1. Effects of CPF on the expression on BCSC and EMT markers in TNBC MDA-MB-231 cells

BCSC clonal expansion and EMT processes are closely related to cancer progression. With the purpose of determining whether CPF modifies TNBC stemness, stemness markers were studied by RT-qPCR in MDA-MB-231 monolayer cells and mammospheres.

In monolayer-grown cells, CPF at 0.05 and 50 μM decreased CD24 (0.75±0.11 and 0.7±0.09-fold below control, *p*<0.01) and ALDH1A1 expression (0.64±0.1, fold below control, *p*<0.05 and 0.47±0.08-fold below control, *p*<0.01; *p*<0.05, respectively). CPF 50 μM also increased CD44 expression (11.84±1.4-fold above control, (*p*<0.01) (Fig. 1A).

**Fig. 1.**
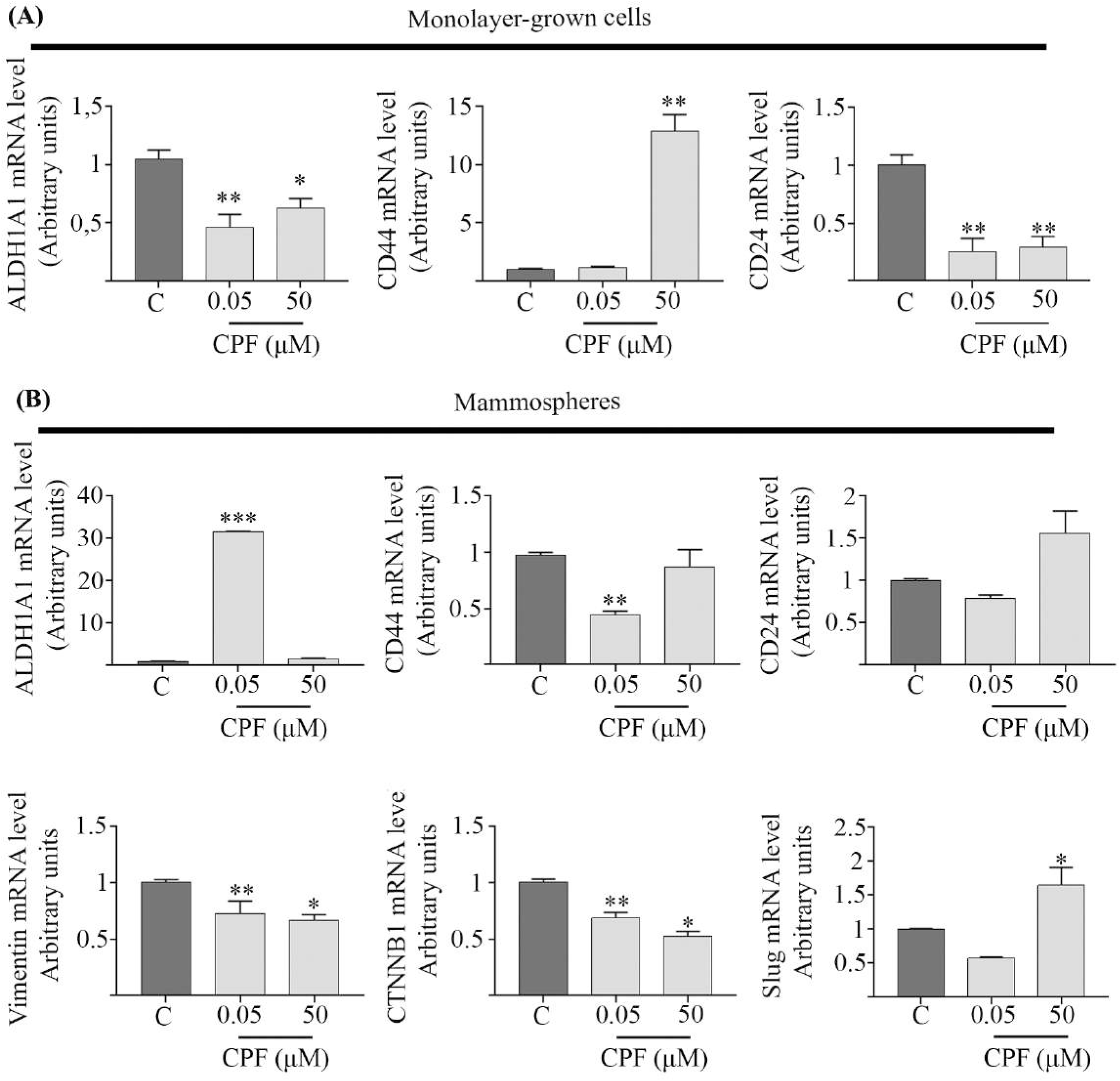
CPF modulates the expression of BCSC and EMT markers in TNBC MDA-MB-231 cells. **(A)–** MDA-MB-231 cells grown in monolayer were exposed to CPF (0.05 and 50 μM) or vehicle (C) for 72h. Relative mRNA levels of ALDH1A1, CD44, CD24 mRNA were determined by RT-qPCR. **(B)–** Relative mRNA levels of ALDH1A1, CD44, CD24, Vimentin, β-catenin and Slug mRNA were determined by RT-qPCR in the mammospheres obtained from MDA-MB-231 cells previously treated for 72 hours with CPF (0.05 and 50 μM) or vehicle (C). Our results were normalized with respect to β-actin mRNA expression. Graphs show the results expressed as mean values ± SEM. One-factor ANOVA and Dunnett’s *a posteriori* test (\**p*<0.05; \*\**p*<0.01; \*\*\**p*<0.001 vs C).

In mammospheres obtained from MDA-MB-231 cells previously exposed to CPF at 0.05 or 50 μM for 72 h, we found that CPF 0.05 μM, diminished CD44 mRNA expression (0.56 ± 0.03-fold below control, *p*<0.05), but a significant increase in ALDH1A1 mRNA expression was induced (30.65 ± 0.16-fold above control, *p*<0.001) (Fig. 1B).

Furthermore, we analysed EMT markers of Vimentin, β-catenin, and Slug mRNA expression in mammospheres obtained from cells previously exposed for 72 h to CPF. We found that CPF 0.05 μM decreased Vimentin (0.38 ± 0.01-fold above control, *p*<0.01), Slug (0.43 ± 0.02-fold above control, *p*=ns) and β-catenin (0.54 ± 0.07-fold above control, *p*<0.01) mRNA expression. In mammospheres obtained from cells exposed to 50 μM CPF we observed a decrease in Vimentin mRNA expression (0.33 ± 0.04-fold below control, *p*<0.01) and β-catenin (0.37 ± 0.08-fold below control, *p*<0.05). In addition, we observed an increase in the expression of Slug mRNA (0.64 ± 0.26-fold above control, *p*<0.05).

### 4.2. CPF effects on triple negative tumor development

The effects of CPF were studied in a triple-negative tumour model in mice. In animals exposed to CPF 0.001 mg/kg/day we observed a significant increase in tumour growth from day 11 after treatment (*day 11*: *p*<0.01, *day 13*: *p*<0.001, *day 15*: *p*<0.01, *day 17*: *p*<0.01, *day 20*: *p*<0.05, *day 22*: *p*<0.05, CPF 0.001 mg/kg/day vs Control). In the group of animals exposed to CPF 0.1 mg/kg/day the significant increase in tumour size was observed later, from day 20 (*day 20*: *p*<0.01, *day 22*: *p*<0.01; CPF 0.1 mg/kg/day vs Control) (Fig. 2A). Parallelly, tumor doubling time in CPF-exposed animals was significantly shorter (7.38 ± 1.34 days for CPF 0.001 mg/kg/day, *p*<0.05 and 6.49 ± 0.91 days for CPF 0.1 mg/kg/day, *p*<0.01) than control (9.49 ± 1.34 days) (Fig. 2B). Fig. 2.C shows the final relative tumor volume for each group. It was found a greater relative tumor volume in the animals treated with CPF, since this was 6.77 ± 0.18 for the control group, 10.59 ± 1.32 for animals exposed to CPF 0.001 mg/kg/day and 9.9 ± 0.35 for the group exposed to CPF 0.1 mg/kg/day, *p*<0.05. In concordance, we observed that both concentrations of CPF significantly increased the percentage of PCNA positive cells in the tumor tissue (control: 26.75 ± 2.87 %, CPF 0.001: 41 ± 3.24%, CPF 0.1: 46.15 ± 3.13%, *p*<0.05 vs control) as shown in Fig. 2D.

**Fig. 2.**
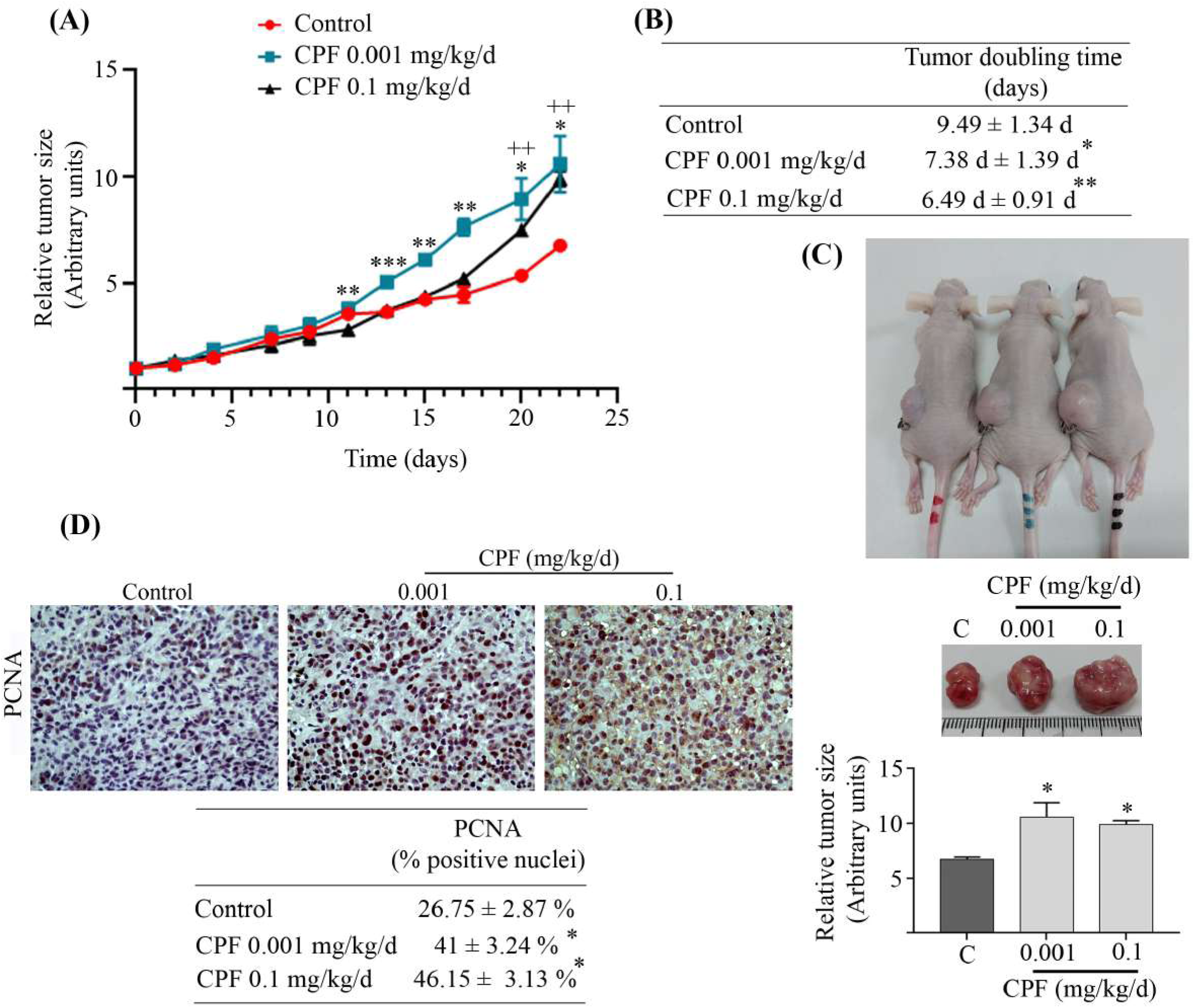
CPF induces an increase in tumor size and cell proliferation. **(A)–** The graph shows the relative tumor volume as a function of time. This parameter was adjusted using a non-linear regression program. Mean value ± SEM of two experiments is represented; *n*=3. Two-factor ANOVA test and Tukey’s a posteriori test (\**p*<0.05, \*\**p*<0.01, \*\*\**p*<0.001 CPF 0.001 vs Control; ^+^*p*<0.05, ^++^*p*<0.01 CPF 0.1 vs control). **(B)–** Tumor doubling time was interpolated from tumor growth curves. The mean ± SEM of the doubling times for the control and CPF-treated groups are shown in the table. One-factor ANOVA test and *a posteriori* Dunnett’s test (\**p*<0.05, \*\**p*<0.01 vs control). **(C)–** Determination of relative tumor volume. Mean ± SEM of two experiments is represented: *n*=3. One– factor ANOVA test and Dunnett’s a posteriori test (\**p*<0.05). **(D)–** PCNA expression was evaluated by immunohistochemistry in tumor. PCNA-positive nuclei are in brown. Magnification: 400X. Percentage of PCNA-positive cells was calculated as the number of positive cells/total number of cells per field. Five randomly selected microscope fields per sample were assessed. Data represent mean valuesJ±JSEM of two experiments is represented *(*p*<0.05; One-way ANOVA).

### 4.3. Evaluation of CPF action on tumor histology, angiogenesis and metastases

Animals in control and exposed groups developed a carcinoma with a solid architectural pattern, accompanied by a marked nuclear pleomorphism, frequent macronuclei and conspicuous nucleoli. However, only animals exposed to CPF had tumors invading the surrounding muscle bundles at the time of euthanasia (Fig. 3A). With TRI, accompanying fibrosis was mild to moderate in the control group and CPF 0.001 mg/kg/day exposed animals, while in the group of animals exposed to CPF 0.1 mg/kg/day, fibrosis was mild to minimal (Fig. 3A). The mitotic rate was found to be significantly higher in animals exposed to CPF 0.1 mg/kg/day with 101.5 ± 18.5 mitoses per 10 high power field (HPF) compared to control 31.3 ± 24 mitoses, *p<*0.05 (Fig. 3B). The mitotic figures were aberrant (Fig. 3A). In assessing necrosis, we found a lower percentage of necrosis in the 0.1 mg/kg/day group compared to the control (32.5 ± 12.5 vs. 68.33 ± 19.2, *p*<0.05) (Fig. 3C). However, the group of animals treated with CPF 0.001 mg/kg/day did not show significant differences in the percentage of necrosis compared to the control group (58.33 ± 9.61 vs. 68.33 ± 19.2%).

**Fig. 3.**
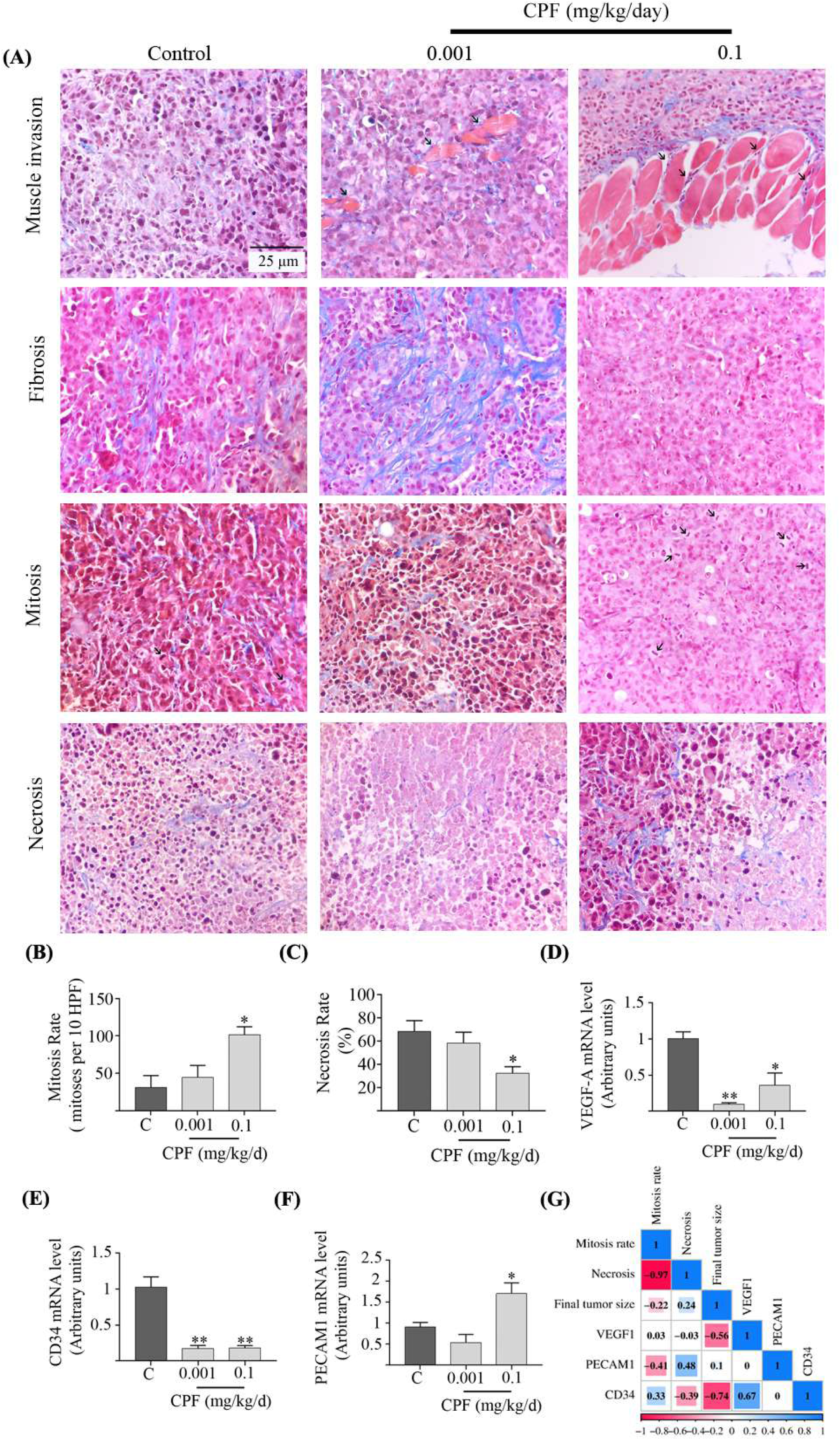
CPF is linked to infiltration of muscle fibers adjacent to the tumor and modulates the expression of angiogenic factors. **(A)–** Representative photographs of tumor characteristics: *First line*: muscle tumor invasion; the black arrows point to the tumor infiltration of the muscle. *Second line*: tumor fibrosis*. Third line:* mitotic cells, marked with black arrows. *Fourth line*: tumor necrosis. Slides were stained with TRI coloring and the photographs were captured with magnification X400; scale bar 25 μm. **(B)–** The mitosis rate (mitoses per 10 HPF) in tumor tissue is shown. Mean value ± SEM of two experiments is represented; n=3. One-factor ANOVA test and Dunnett’s a posteriori test (*p<0.05). **(C)–** The percentage of necrotic area in tumor tissue is shown. Mean value ± SEM of two experiments is represented; *n*=3. One-factor ANOVA test and Dunnett’s a posteriori test (\**p*<0.05). Relative levels of VEGF-A **(D)**, CD34 **(E)** and PECAM1 **(F)** mRNA expression in tumors were determined by RT-qPCR. The results were normalized with respect to β-actin mRNA expression. The graphs show the results expressed as the mean values ± SEM. One-factor ANOVA test and Dunnett’s *a posteriori* test (\**p*<0,05; \*\**p*<0,01; vs C). **(G)–**Correlation matrix plot with lower triangle color intensity and size of the square proportional to the correlation coefficients. The color scale bar ranged from −1.0 (red) to 1.0 (blue). The size and color of the circles indicate the magnitude of the correlation between parameters. Red and blue, respectively, denote negative and positive correlations. These were performed in SRplot (Tang et al., 2023).

The expression of markers related to tumor angiogenesis showed that CPF downregulates VEGF-A (CPF 0.001 mg/kg/day: 0.9 ± 0.02-fold above control, *p*<0.01; CPF 0.1 mg/kg/day: 0.64 ± 0.16-fold above control, *p*<0.05) and CD34 (0.83 ± 0.04-fold above control, *p*<0.01; 0.82 ± 0.03-fold above control, *p*<0.01, respectively) mRNA level (Fig 3D-E). Additionally, we found an increase in PECAM1 mRNA expression in the group exposed to 0.1 mg/kg/day (0.79 ± 0.25-fold over control, *p*<0.05) (Fig. 3F).

When analysing the correlation among the above-mentioned factors (Fig. 3G), we observed a negative correlation between final tumour volume and VEGF1 mRNA level (*r^2^*= –0.56, *p*<0.02) as well as between final tumour volume and CD34 (*r^2^*= –0.74, *p*<0.001). Likewise, this was found between necrosis and mitotic rate (*r^2^*= –0.97, *p*<0.001), and necrosis and PECAM1 mRNA level (*r^2^*= –0.41, *p*<0.08). Lastly, there was a positive correlation between mitotic rate and PECAM1 (*r^2^*= 0.48, *p*<0.04) and between VEGF1 and CD34 mRNA levels (*r^2^*= 0.67, p<0.001).

We found lung metastases in both the animals from the control group and in those exposed to CPF 0.001 mg/kg/day, but the metastatic areas were greater in the latter, as shown in Fig. 4. In contrast, the animals treated with CPF 0.1 mg/kg/day had no metastases but showed a marked disruption of the normal structure of lung tissue due to abundant interstitial inflammatory infiltrate and alveolar edema.

**Fig. 4.**
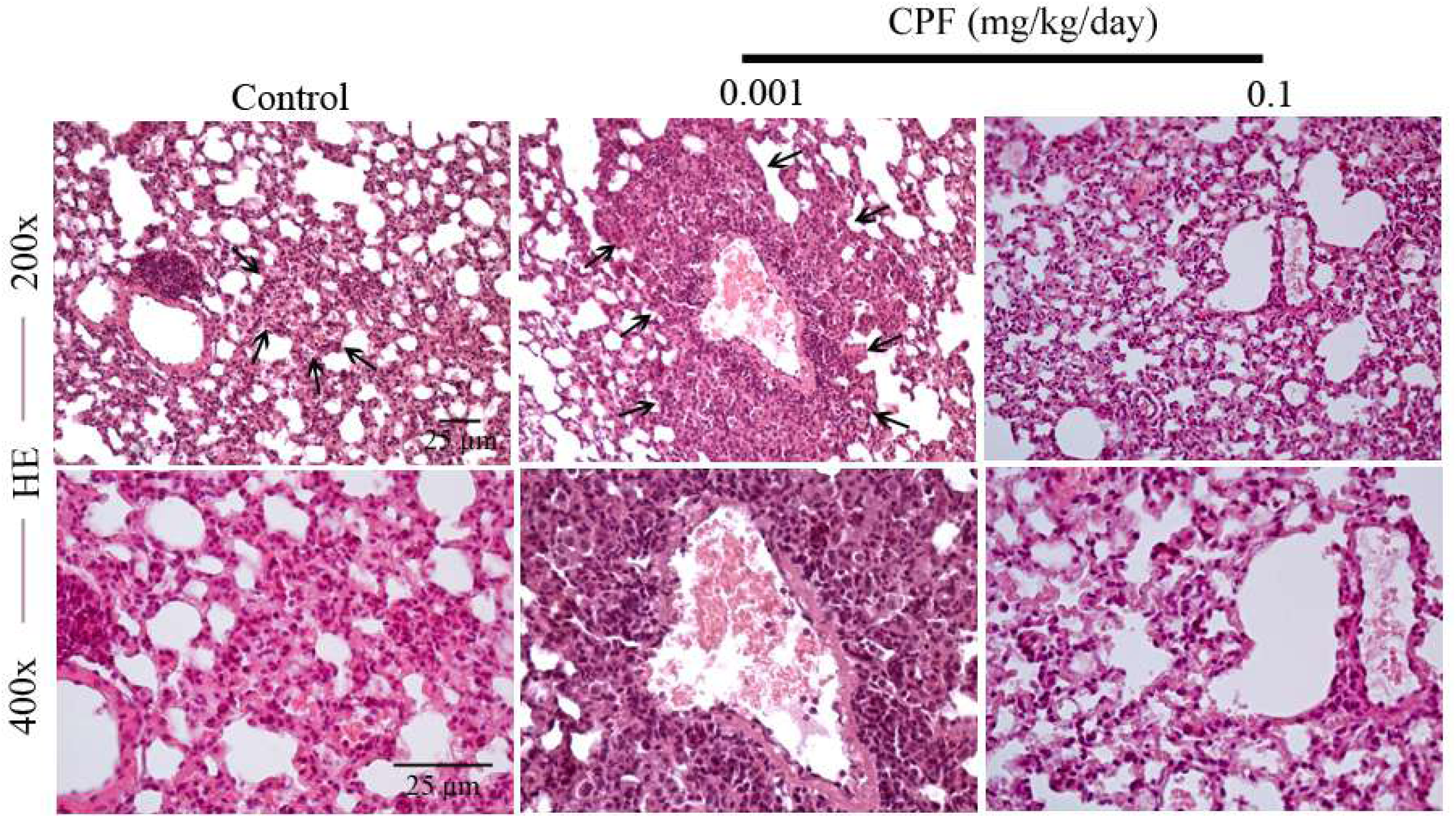
Low dose of CPF induces lung metastases. Representative photographs illustrating histology of HE-stained lung tissue. Black arrows point to regions showing lung metastases. Scale bar: 25 μm. Magnification 200 and 400x.

### 4.4 Effects of CPF on the expression of BCSC and EMT markers in tumor tissue

We determined the expression of BCSC markers in the tumors of the animals exposed or not to CPF (Fig. 5). We found an increase in ALDH1A1 mRNA levels in CPF 0.001 mg/kg/day group (1.18 ± 0.04-fold over control, *p*<0.001) and CPF 0.1 mg/kg/day group (1.33 ± 0.08-fold over control, *p*<0,001). Moreover, in the CPF 0.1 mg/kg/day group we observed an upregulation of CD44 (0.7 ± 0.10-fold over control, *p*<0.001) and CD24 (17.89 ± 1.73-fold over control, *p*<0.01) markers. In addition, we found a significant increase in mRNA expression of Slug (17.81 ± 3.0-fold over control, *p*<0.01) and Vimentin levels (0.75 ± 0.10-fold over control, *p*<0.001) and a decrease in CTNNB1 (0.66 ± 0.05-fold over control, *p*<0.05) in the CPF 0.1 group.

**Fig. 5.**
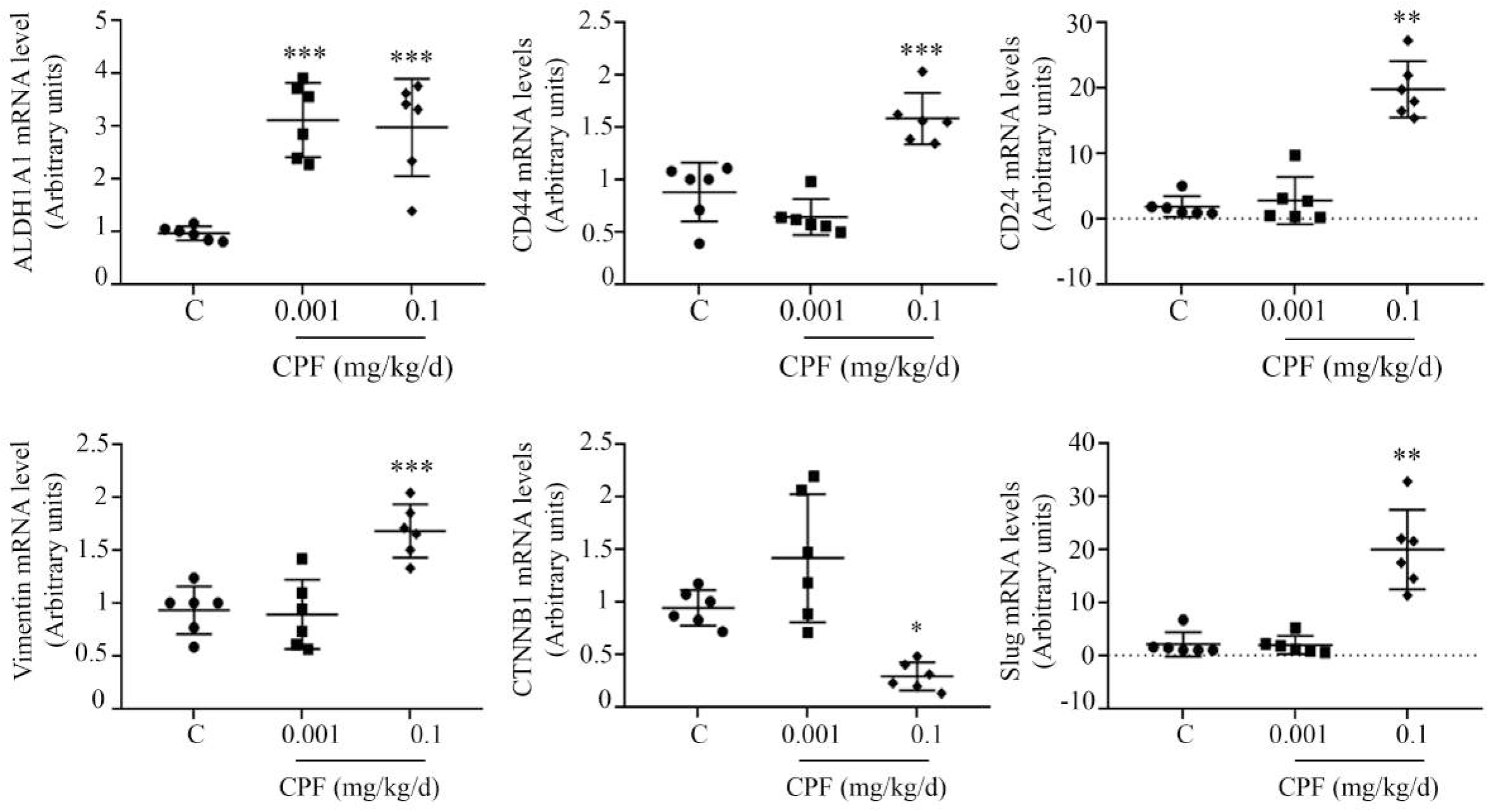
CPF regulates mRNA expression of BCSC and EMT markers in tumor tissue. Relative levels of ALDH1A1, CD44, CD24, Vimentin, CTNNB1 and Slug mRNA expression in tumors were determined by RT-qPCR. The results were normalized with respect to β-actin mRNA expression. The graphs show the results expressed as the mean values ± SEM. One-factor ANOVA test and Dunnett’s a posteriori test (\**p*<0,05; \*\**p*<0,01; \*\*\**p*<0.001 vs C).

In addition, we analyzed the correlation between EMT and BCSC markers (Fig. 6A) in the tumors of animals from each group. We found a positive correlation between Vimentin and Slug mRNA levels with CD44 (*r^2^*=0.634, *p*=0.005; *r^2^*=0.840, *p*=0.001, respectively) and CD24 (*r^2^*=0.830, *p*=0.001; *r^2^*=0.863, *p*=0.001, respectively), while the correlation with CTNNB1 was negative for both BCSC markers (CD44: *r^2^*= –0.768, *p*=0.002; CD24: *r^2^*=– 0.706, *p*=0.001). ALDH1A1 showed no correlation with Vimentin or CTNNB1 levels. We evaluated the correlation of BCSC markers (Fig. 6B) and we found a positive correlation between CD44 and CD24 (*r^2^*=0.754, *p*=0.001). We also found a negative relationship of CTNNB1 with Slug and Vimentin (*r^2^*= –0.688, *p*=0.0016; *r^2^*=-0.744, *p*=0.0004, respectively), while the correlation was positive between the latter two markers (*r^2^*=0.649, *p*=0.0035) as shown in Figure 6C.

**Fig. 6.**
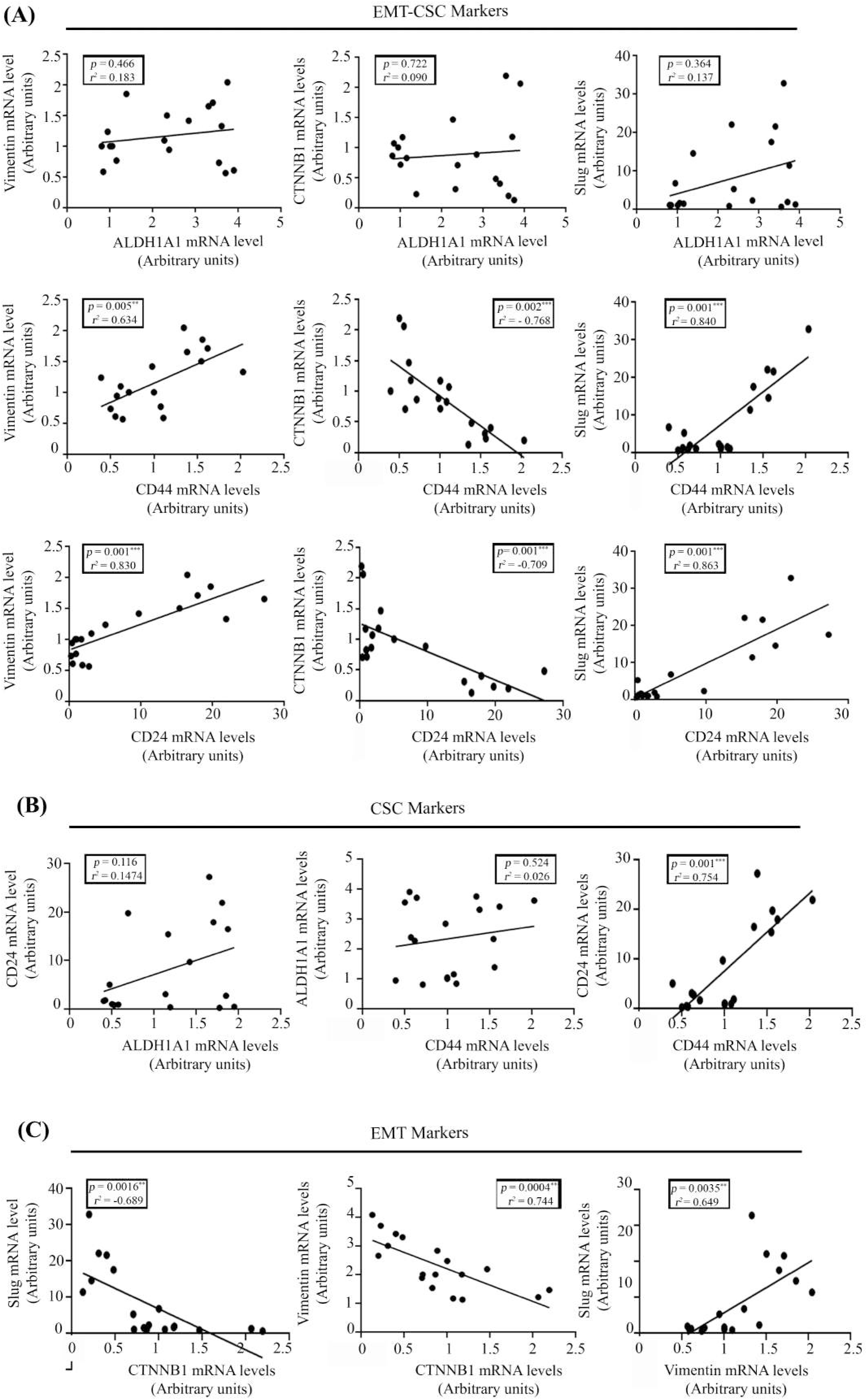
Strong correlation between EMT and BCSC markers in CPF-exposed animals. **(A)–** ALDH1A1, CD44 and CD24 mRNA levels were correlated with Vimentin, CTNNB1 and Slug. **(B)–** ALDH1A1, CD44 and CD24 mRNA levels were correlated with each other. **(C)** CTNNB1, Slug and Vimentin mRNA levels were correlated with each other. Pearson’s test was applied. Values of *r^2^* greater than 0.7 (or less than –0.7) will be considered as a strong correlation. (\*\**p*<0.01; \*\*\**p*<0.001).

Finally, we explored the expression of β-catenin, Slug and CD44 in the tumors (Fig. 7). We found increased expression and nuclear translocation of β-catenin in animals treated with CPF 0.001 mg/kg/day and an abolition of expression in those neoplasms developed in animals exposed to CPF 0.1 mg/kg/day. Furthermore, we observed an increase in Slug expression in tumors grown in both groups of animals exposed to CPF and an increase in nuclear translocation in the group exposed to the high dose. In exposed mice, we observed an increase in the expression of CD44 and the localization of this protein predominantly in the plasma membrane, where it functions as a co-receptor for a wide diversity of extracellular matrix ligands, such as hyaluronic acid, being able to activate diverse signalling pathways (Al-Othman et al., 2020).

**Fig. 7.**
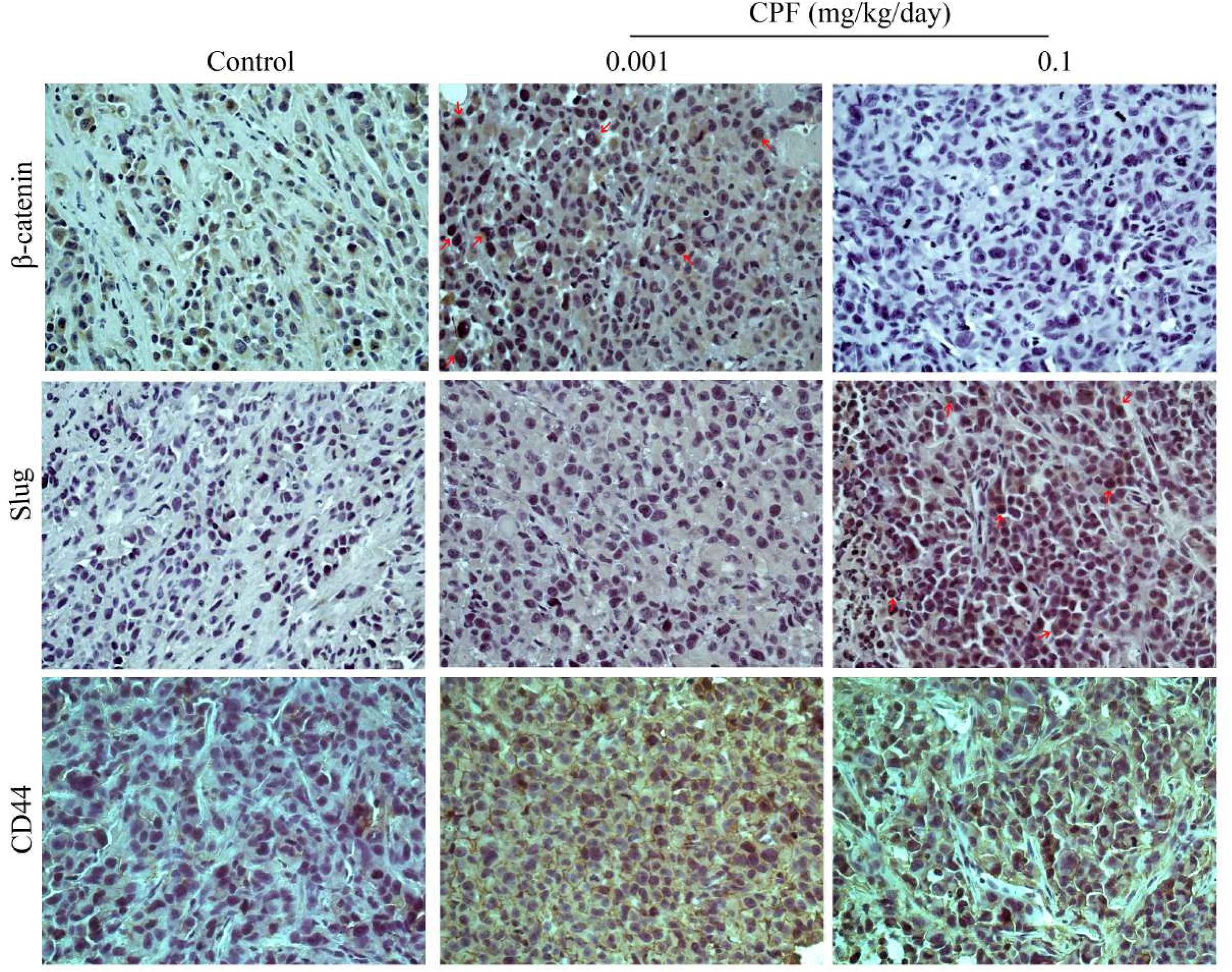
CPF modulates protein expression and subcellular localization of BCSC and EMT markers in tumor tissue. β-catenin, Slug and CD44 protein expression and subcellular localization were determined in tumor tissue by immunohistochemical technique (IHC). Representative photographs corresponding to the control, CPF 0.001 and CPF 0.1 mg/kg/day experimental groups are shown. Red arrows show the nuclear translocation of Slug and β-catenin. Scale bar: 25 μm. Magnification 400x.

## 5. Discussion

Breast cancer is a prevalent human malignancy and a very common cause of cancer– related death among women worldwide. It is currently considered a multifactorial disease, and environmental, hormonal, genetic, lifestyle and nutritional influences are implicated in its genesis (Palomeras et al., 2018). TNBC is a type of breast cancer that affects 15–20% of the patients. It is regarded as an aggressive tumor with a faster growth rate and worse prognosis than estrogen receptor-positive breast cancer (Bauer et al., 2007).

Despite advances in the diagnosis and treatment of breast cancer, many patients fail therapy, leading to disease progression, recurrence and reduced overall survival. There is growing evidence proposing that tumors are composed of heterogeneous populations of cells with a hierarchical organization. Among these, BCSC often have a higher tolerability to chemotherapy, hormone therapy and radiotherapy, and can reproduce the tumor after depletion of cell populations sensitive to first-line therapy, leading to disease relapse.

In this study, we demonstrated that CPF can modulate the expression of EMT and BCSC markers in *in vitro* and *in vivo* models. In the *in vivo* triple negative tumor model, CPF decreased the doubling period increasing the primary tumor volume. Furthermore, CPF augmented cell proliferation, mitotic rate and stimulated the infiltration of adjacent muscle fibers. In addition, in the group of animals exposed to the low dose of the pesticide, an increase in the size and number of lung metastases were found. All our studies were performed with CPF concentrations on the order of those that can be orally incorporated through food and water intake and support our hypothesis that CPF is a risk factor for BC. Dose selection was justified in the MM section and discussed in previous publications of our group (Lasagna et al., 2020, 2022; Ramos Nieto et al., 2021; Ventura et al., 2016).

Since that CD44, CD24 and ALDH1A1 were found expressed in BCSC from a wide variety of carcinomas, including pancreatic, colon, lung, ovarian, prostate and breast (Eramo et al., 2008; Thiery et al., 2009), we evaluated those BCSC markers in cells grown in monolayer and tridimensional mammospheres cultures derived from CPF-exposed TNBC cells. These results clearly support the role of CPF on breast cancer cell transformation.

We found that CPF modulates the plasticity of BCSCs since it promotes epithelial to mesenchymal transformation depending on the microenvironment in which the exposure takes place. The plasticity of BCSC plays a crucial role in the ability of these cells to generate distant metastases (Liu et al., 2014).

In this sense, we found that CPF induces EMT-like BCSC when the cells were grown in monolayer cultures. This mesenchymal phenotype was manifested by the presence of CD24^-^/CD44^+^ cells. However, a significant reduction of ALDH1A1 expression was detected. Our experiments performed in the mammospheres obtained from cells previously exposed to CPF showed a mesenchymal epithelial transition-like BCSC phenotype, characterized by increased expression of ALDH1A1, accompanied by down-regulation of CD44, Vimentin and β-catenin. The epithelial phenotype has been associated with expression of epithelial markers, down-regulation of mesenchymal markers and presence of cell polarity. This cellular subpopulation is usually found more localized in the center of tumors and exhibits intense proliferation (Liu et al., 2014).

In the *in vivo* TNBC model, we determined that CPF generated an increase in final tumor volume and a decrease in doubling time, along with an increment of PCNA positive cells in the tumor. It is widely known that the tumor growth is usually accompanied by the generation of hypoxic regions. Despite developing larger tumors, the animals treated with CPF 0.001 mg/kg/day showed a percentage of necrosis similar to the control group. In opposition, animals treated with CPF 0.1 mg/kg/day showed significantly smaller areas of necrosis than controls, with a high rate of mitosis. Hypoxia is prominent in the microenvironment of breast tumors and plays a central role in regulating stemness phenotype and function (Conley et al., 2012). This condition generates an increase in this BCSC, enhances their self-renewal capacity and promotes the maintenance in their undifferentiated state. All these processes are commonly involved in tumor progression and metastases.

Although angiogenesis has been a long-standing therapeutic target in malignant tumors, we observed that CPF would seem to exhibit antiangiogenic effects because it down– regulates VEGF-A and CD34 mRNA levels in CPF treated groups. We have also detected an increase of proliferation of TNBC cells in these groups with respect to control. It could be explained by different survival mechanisms carried out by tumor cells. Warburg effect is one of the known processes described when hypoxia condition must be overcome. This effect has been defined as aerobic glycolysis and is essential for rapid cell division and survival (de Heer et al., 2020). Furthermore, in animals exposed to the high dose of CPF we found an up-regulation in PECAM-1 mRNA expression. This marker correlates positively with mitotic rate and negatively with necrosis. These results are consistent with those reported by Abraham et al. (2018), who found that PECAM-1 is involved in modulating the tumour microenvironment and promoting tumour cell proliferation, rather than by stimulating tumour angiogenesis.

In our experiments, the animals exposed to CPF 0.001 mg/kg/day developed larger lung metastases. Coincidentally, the tumors from the same group were able to infiltrate adjacent muscle fibers which may be associated with high aggressiveness and local progression. Metastasis is a complex process involving the spread of malignant cells from a primary tumour to distal organs. To achieve this, tumor cells must exhibit phenotypic plasticity in both directions. Firstly, they must undergo EMT to escape from the primary tumor and enter the bloodstream. Conversely, mesenchymal-epithelial transition often occurs to colonize the metastatic site (Ionkina et al., 2021). Therefore, although there are multiple *in vivo* experimental models of metastasis, we believe that xenotransplantation in immunocompromised mice reproduces the entire metastatic cascade unlike models in which cells are inoculated directly into the tail vein, bypassing many of the aforementioned stages (Tuomela and Härkönen, 2014). In contrast, no metastases were observed in animals exposed to CPF 0.1 mg/kg/day, but a significant inflammatory infiltrate and loss of lung histoarchitecture were detected. It was reported that very high doses of CPF can alter lung structure due to its toxicity mediated by an imbalance of oxido-reduction mechanisms and increased proinflammatory cytokines (Alipanah et al., 2022).

Furthermore, both doses of CPF enhanced ALDH1A1 mRNA expression in tumors. Also, the higher dose of CPF up-regulated CD44 and CD24 expression. Ginestier et al., (2007) reported that the upregulation of these markers are indicative of greatest tumor-initiating capacity. In their work, they observed that the injection of a few cells co-expressing high levels of CD24, CD44 and ALDH1A1 were enough to generate *de novo* tumors in mice.

ALDH1 is one of the most common markers of BCSC, and the ALDH1A1 isoenzyme regulates the expression of several genes involved in stemness maintenance and self– renewal promoting tumor growth and drug resistance (Ciccone et al., 2018; Vassalli, 2019). High ALDH1A1 expression was associated with poor prognostic features, including high– grade Nottingham Prognostic Index, lymph node metastases, and highly proliferative subtypes such as ER+ (luminal B) and TNBC. The expression of ALDH1A1 was positively correlated with the expression of CD44, CD24, TWIST, SOX9, EPCAM, and CD133 (Althobiti et al., 2020).

In the group of animals exposed to CPF 0.1 mg/kg/day we observed an up-regulation of CD44 and CD24 accompanying the increased expression of ALDH1A1. Additionally, we found a positive correlation between CD44 and CD24 with each other, as well as with the EMT markers Vimentin and Slug. Strikingly, CD44 and CD24 correlated negatively with β– Catenin. This result is supported by a paper published by Jang et al, 2015, in which they reported that some EMT markers, such as Vimentin, correlated positively with those cells with CD44^+^/CD24^-^ phenotype, while tumor cells expressing ALDH1A1 showed no correlation with these markers (Jang et al., 2015). Furthermore, CD44 mRNA levels were inversely correlated with ER, PR and HER2 status, suggesting that it often represents a marker of basal-like tumors. These results are in the same direction as our findings in the present study. Additionally, Xu et al. (2016) found that CD44 overexpression in patients with basal-like breast cancer is positively associated with decreased overall survival, metastases-free survival, and recurrence. Thus, the authors proposed that CD44 could be a potential prognostic marker of breast cancer and, therefore, may serve as a therapeutic target for the basal subtype.

Surprisingly, CD24 was elevated in animals exposed to the high dose of CPF. Nevertheless, some evidences indicate that Slug overexpression, could lead to a switch from CD44^+^/CD24^-^ to CD44^+^/CD24^+^ cells, which retain the ability to induce mammosphere formation and stemness (Bhat-Nakshatri et al., 2010). Additionally, the CD24-P-Selectin pathway promotes the interaction of tumor cells with endothelial cells and platelets, enhancing metastatic dissemination and leading to the activation of the c-SRC pathway, which could be the starting point of many other signaling pathways such as AKT and Focal Adhesion Kinase (Altevogt et al., 2021). Indeed, this is consistent with our previous reports, where we demonstrated that CPF induced c-SRC activation and MMP2 expression and activity, reduced β-catenin, increased Vimentin expression and migration (Lasagna et al., 2020).

Lastly, we evaluated the modulation of Slug, β-catenin and CD44 expression and their localization. A striking finding was the opposing effect generated by both CPF doses on the expression of β-catenin, which represents the central component of the canonical Wnt signaling pathway. This protein binds to the cytoplasmic tail of E-cadherin to maintain cell– cell adhesion (Xu, Zhang, Xu, & Jiang, 2020). E-cadherin is frequently mutated or silenced in TNBC, leading to the release of β-catenin into the cytoplasm (Liu et al., 2013). The concentration of cytoplasmic β-catenin is usually carefully controlled by the destruction complex, but the components of this complex are frequently mutated, deleted, hypermethylated or reduced in breast cancer, which increases the stability of cytoplasmic β-Catenin and the likelihood of β-catenin entering the nucleus. This effect was observed in animals exposed to the low dose of CPF. Moreover, non-canonical Wnt signaling is preferentially activated in TNBC and is mainly induced by epigenetic activation of non– canonical Wnts and their receptors (Xu et al., 2020). These pathways could be activated by the high concentration of CPF and explain, at least in part, why we observed a reduction in both β-catenin and CTNNB1 gene expression in tumors from animals exposed to this dose.

We showed that the pesticide CPF at both doses enhanced Slug expression. The high dose also increased its nuclear translocation. Slug is a member of the SNAIL superfamily of zinc-finger transcription factors and plays a critical role in modulating EMT during carcinogenesis (Georgakopoulos-Soares et al., 2020). It is known that Slug not only acts as a transcriptional repressor factor to block the expression of epithelial markers (e.g., E– cadherin, occludin, and Claudin-1), but also directly activates the expression of ZEB1 (Wels et al., 2011), an inducer of EMT. A recent meta-analysis (Zhang et al., 2022) demonstrated that Slug protein overexpression resulted in a worse overall survival and disease-free survival in breast cancer patients. Consequently, the up regulation of Slug at the tumor level is further evidence of the ability of CPF to induce EMT in an *in vivo* model and thus promote the development of distant metastases.

## 6. Conclusions

All the above findings confirm that CPF is able to promote breast cancer progression through modulation of BCSC and EMT marker expression and generate lung metastases in an *in vivo* model. Fortunately, a progressive elimination of CPF-containing formulations was established in Argentina in 2021 until their total ban for agricultural use on all crops. Despite this, it is necessary to strengthen control measures to ensure compliance with this regulation, to prevent general population from the very harmful health effects of continuous exposure to this toxic substance.

---

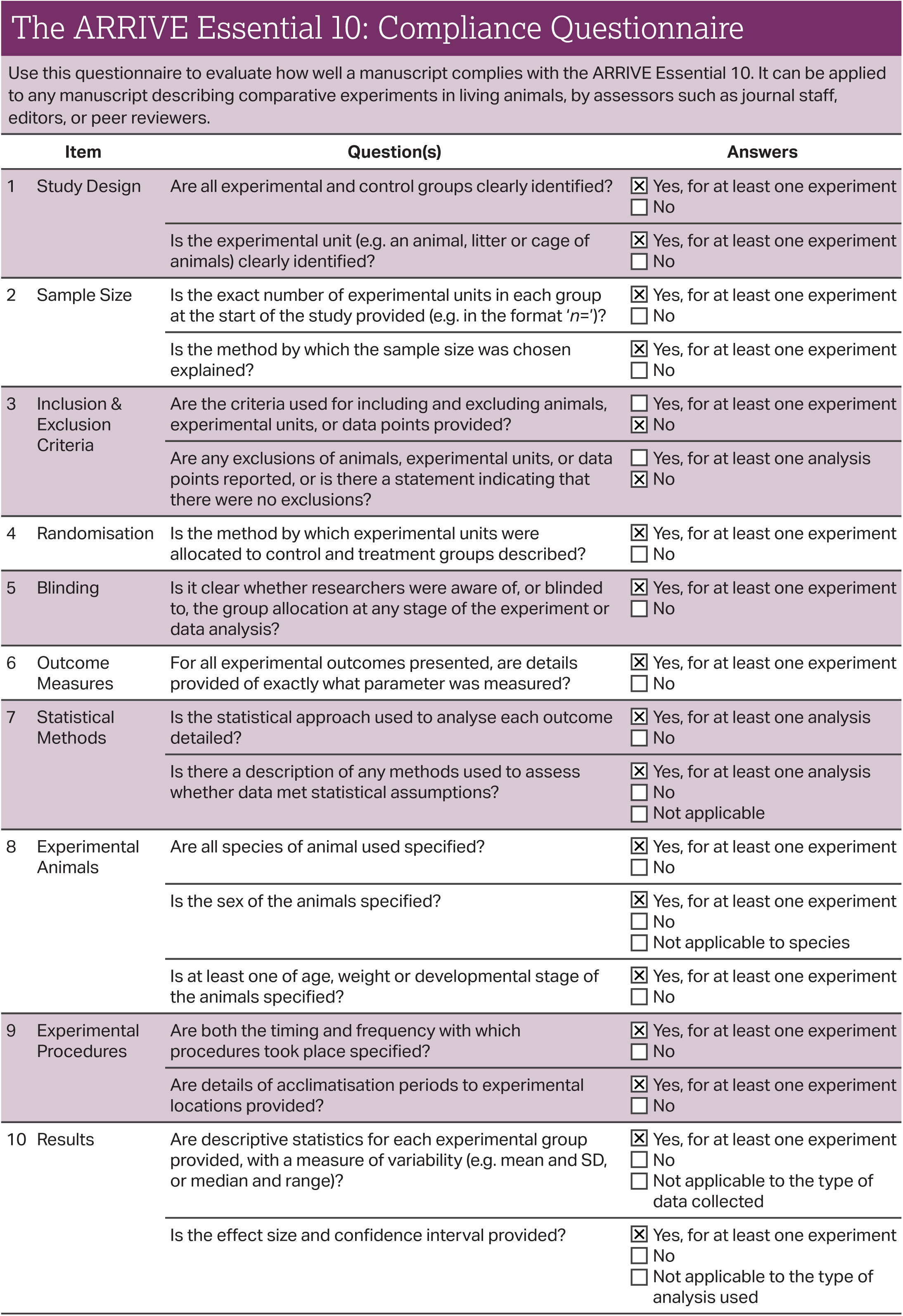

## Notes on questionnaire design

The ARRIVE guidelines are a useful resource for authors preparing manuscripts describing animal research, and also provide a framework to evaluate the transparency of those manuscripts. To assess reporting quality, numerous studies have in the past sought to operationalise reporting guidelines (including ARRIVE). Typically, this involves scoring a manuscript’s degree of compliance with guideline items in a binary fashion (e.g. an item is either not reported or reported) [1-3], a graded fashion (e.g. not, partially, or completely reported) [4,5], or a combination of the two [6].

This questionnaire has been designed to be as concise and user-friendly as possible. The number of questions used to assess a manuscript’s compliance has been kept to a minimum, and in most cases each question is designed to be answered in a binary fashion. Compliance with some Essential 10 sub-items is inherently impossible to judge in this way, instead requiring a subjective judgement on the level of detail provided. For this reason, not all sub-items are represented by a question in this questionnaire.

To facilitate binary answers, it has been necessary to identify the minimum information in a manuscript sufficient to comply with each question. The strengths of this approach include the relatively short length of the questionnaire (and the correspondingly low time burden of using it), and the avoidance of ambiguity that would arise from a graded answering system, in which an intermediate score (e.g. ‘partially/insufficiently reported’) could denote a number of distinct deficiencies in compliance with an item (e.g. either only part of the item was complied with, or only the reporting of some experiments in the manuscript complied with the item.)

Limitations of this approach centre on the necessity to identify the minimum information sufficient to comply with each question. In some cases, this has resulted in questions that require a guideline sub-item’s criteria to have been fulfilled in the reporting of only one experiment in a manuscript. As a result, not all experiments in a manuscript may be described in a way that fulfils that criterion, despite the manuscript being considered to comply with the guidelines overall.

---

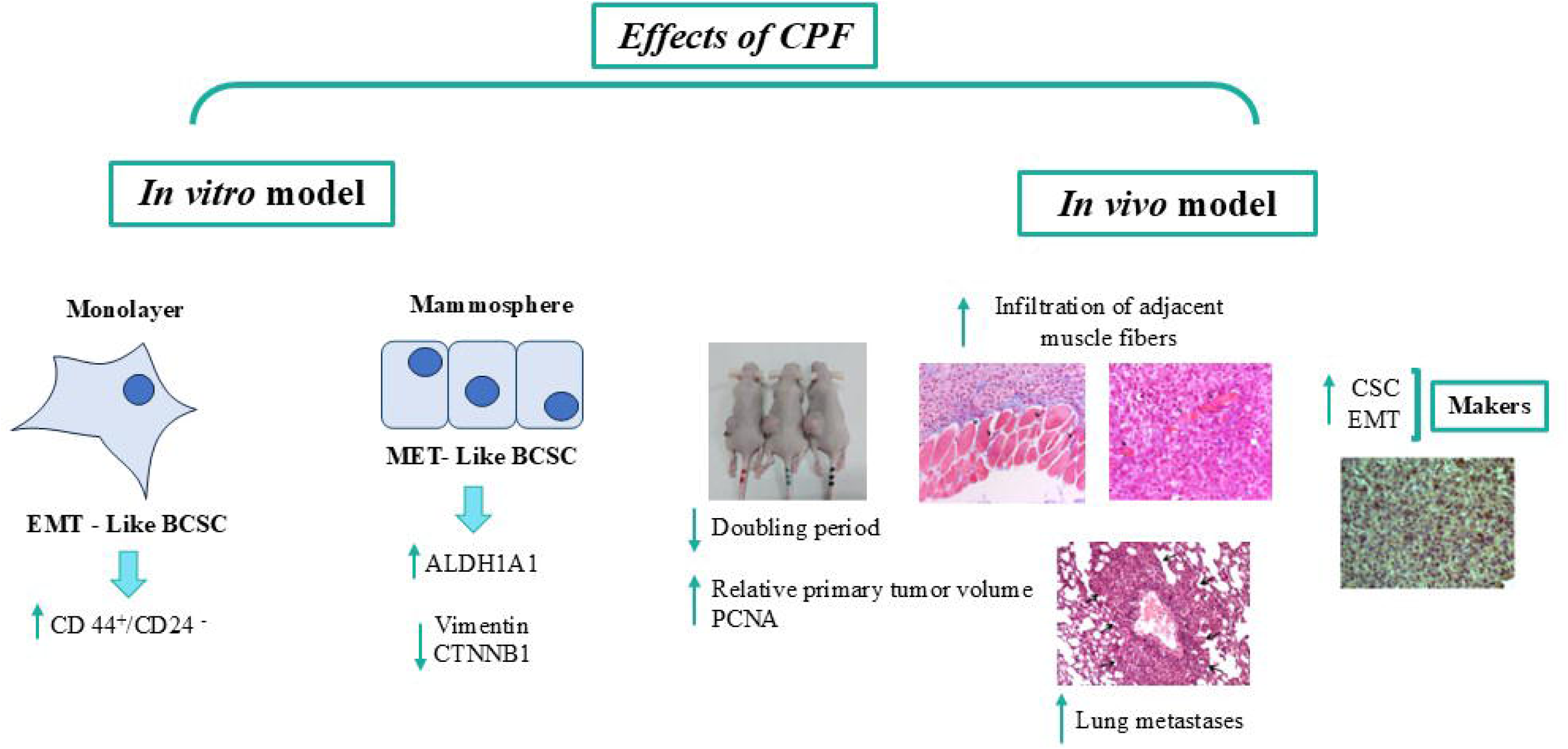

## References

1. Abraham, V., Cao, G., Parambath, A., Lawal, F., Handumrongkul, C., Debs, R., DeLisser, H.M., 2018. Involvement of TIMP-1 in PECAM-1-mediated tumor dissemination. Int. J. Oncol. 53, 488–502. 10.3892/IJO.2018.4422

2. Al-Hajj, M., Wicha, M.S., Benito-Hernandez, A., Morrison, S.J., Clarke, M.F., 2003. Prospective identification of tumorigenic breast cancer cells. Proc. Natl. Acad. Sci. U. S. A. 100, 3983. 10.1073/PNAS.0530291100

3. Al-Othman, N., Alhendi, A., Ihbaisha, M., Barahmeh, M., Alqaraleh, M., Al-Momany, B.Z., 2020. Role of CD44 in breast cancer. Breast Dis. 39, 1–13. 10.3233/BD-190409

4. Alipanah, H., Kabi Doraghi, H., Sayadi, M., Nematollahi, A., Soltani Hekmat, A., Nejati, R., 2022. Subacute toxicity of chlorpyrifos on histopathological damages, antioxidant activity, and pro-inflammatory cytokines in the rat model. Environ. Toxicol. 37, 880– 888. 10.1002/TOX.23451

5. Altevogt, P., Sammar, M., Hüser, L., Kristiansen, G., 2021. Novel insights into the function of CD24: A driving force in cancer. Int. J. cancer 148, 546–559. 10.1002/IJC.33249

6. Althobiti, M., El Ansari, R., Aleskandarany, M., Joseph, C., Toss, M.S., Green, A.R., Rakha, E.A., 2020. The prognostic significance of ALDH1A1 expression in early invasive breast cancer. Histopathology 77, 437–448. 10.1111/HIS.14129

7. Barkal, A.A., Brewer, R.E., Markovic, M., Kowarsky, M., Barkal, S.A., Zaro, B.W., Krishnan, V., Hatakeyama, J., Dorigo, O., Barkal, L.J., Weissman, I.L., 2019. CD24 signalling through macrophage Siglec-10 is a target for cancer immunotherapy. Nature 572, 392–396. 10.1038/S41586-019-1456-0

8. Batlle, E., Clevers, H., 2017. Cancer stem cells revisited. Nat. Med. 23, 1124–1134. 10.1038/NM.4409

9. Bauer, K.R., Brown, M., Cress, R.D., Parise, C.A., Caggiano, V., 2007. Descriptive analysis of estrogen receptor (ER)-negative, progesterone receptor (PR)-negative, and HER2– negative invasive breast cancer, the so-called triple-negative phenotype: a population– based study from the California cancer Registry. Cancer 109, 1721–1728. 10.1002/CNCR.22618

10. Bhat-Nakshatri, P., Appaiah, H., Ballas, C., Pick-Franke, P., Goulet, R., Badve, S., Srour, E.F., Nakshatri, H., 2010. SLUG/SNAI2 and tumor necrosis factor generate breast cells with CD44+/CD24– phenotype. BMC Cancer 10. 10.1186/1471-2407-10-411

11. Chaffer, C.L., Weinberg, R.A., 2011. A perspective on cancer cell metastasis. Science 331, 1559–1564. 10.1126/SCIENCE.1203543

12. Chakraborty, P.K., Scharner, B., Jurasovic, J., Messner, B., Bernhard, D., Thévenod, F., 2010. Chronic cadmium exposure induces transcriptional activation of the Wnt pathway and upregulation of epithelial-to-mesenchymal transition markers in mouse kidney. Toxicol. Lett. 198, 69–76. 10.1016/j.toxlet.2010.05.007

13. Chang, J.C., 2016. Cancer stem cells: Role in tumor growth, recurrence, metastasis, and treatment resistance. Medicine (Baltimore). 95, S20–S25. 10.1097/MD.0000000000004766

14. Chi, Y., Huang, S., Peng, H., Liu, M., Zhao, J., Shao, Z., Wu, J., 2015. Critical role of CDK11p58 in human breast cancer growth and angiogenesis. BMC Cancer 15, 1–10. 10.1186/S12885-015-1698-7/FIGURES/4

15. Ciccone, V., Terzuoli, E., Donnini, S., Giachetti, A., Morbidelli, L., Ziche, M., 2018. Stemness marker ALDH1A1 promotes tumor angiogenesis via retinoic acid/HIF-1α/VEGF signalling in MCF-7 breast cancer cells. J. Exp. Clin. Cancer Res. 37. 10.1186/S13046-018-0975-0

16. Cocca, C., Martin, G., Rivera, E., Davio, C., Cricco, G., Lemos, B., Fitzsimons, C., Gutierrez, A., Levin, E., Levin, R., Croci, M., Bergoc, R.M., 1998. Original Paper An Experimental Model of Diabetes and Cancer in Rats. Eur J Cancer 34.

17. Conley, S.J., Gheordunescu, E., Kakarala, P., Newman, B., Korkaya, H., Heath, A.N., Clouthier, S.G., Wicha, M.S., 2012. Antiangiogenic agents increase breast cancer stem cells via the generation of tumor hypoxia. Proc. Natl. Acad. Sci. U. S. A. 109, 2784–2789. 10.1073/PNAS.1018866109/SUPPL_FILE/PNAS.201018866SI.PDF

18. de Heer, E.C., Jalving, M., Harris, A.L., 2020. HIFs, angiogenesis, and metabolism: elusive enemies in breast cancer. J. Clin. Invest. 130, 5074–5087. 10.1172/JCI137552

19. Ding, S.Z., Yang, Y.X., Li, X.L., Michelli-Rivera, A., Han, S.Y., Wang, L., Pratheeshkumar, P., Wang, X., Lu, J., Yin, Y.Q., Budhraja, A., Hitron, A.J., 2013. Epithelial– mesenchymal transition during oncogenic transformation induced by hexavalent chromium involves reactive oxygen species-dependent mechanism in lung epithelial cells. Toxicol. Appl. Pharmacol. 269, 61–71. 10.1016/j.taap.2013.03.006

20. Eramo, A., Lotti, F., Sette, G., Pilozzi, E., Biffoni, M., Di Virgilio, A., Conticello, C., Ruco, L., Peschle, C., De Maria, R., 2008. Identification and expansion of the tumorigenic lung cancer stem cell population. Cell Death Differ. 15, 504–514. 10.1038/SJ.CDD.4402283

21. European Food Safety Authority, 2019. Statement on the available outcomes of the human health assessment in the context of the pesticides peer review of the active substance chlorpyrifos. EFSA J. 17. 10.2903/J.EFSA.2019.5809

22. Fang, X., Zheng, P., Tang, J., Liu, Y., 2010. CD24: from A to Z. Cell. Mol. Immunol. 7, 100–103. 10.1038/CMI.2009.119

23. Fares, J., Fares, M.Y., Khachfe, H.H., Salhab, H.A., Fares, Y., 2020. Molecular principles of metastasis: a hallmark of cancer revisited. Signal Transduct. Target. Ther. 5. 10.1038/S41392-020-0134-X

24. Georgakopoulos-Soares, I., Chartoumpekis, D. V., Kyriazopoulou, V., Zaravinos, A., 2020. EMT Factors and Metabolic Pathways in Cancer. Front. Oncol. 10. 10.3389/FONC.2020.00499

25. Ginestier, C., Hur, M.H., Charafe-Jauffret, E., Monville, F., Dutcher, J., Brown, M., Jacquemier, J., Viens, P., Kleer, C.G., Liu, S., Schott, A., Hayes, D., Birnbaum, D., Wicha, M.S., Dontu, G., 2007. ALDH1 is a marker of normal and malignant human mammary stem cells and a predictor of poor clinical outcome. Cell Stem Cell 1, 555–567. 10.1016/J.STEM.2007.08.014

26. Housman, G., Byler, S., Heerboth, S., Lapinska, K., Longacre, M., Snyder, N., Sarkar, S., 2014. Drug Resistance in Cancer: An Overview. Cancers (Basel). 6, 1769–1792. 10.3390/cancers6031769

27. Hu, W.Y., Shi, G. Bin, Hu, D.P., Nelles, J.L., Prins, G.S., 2012. Actions of estrogens and endocrine disrupting chemicals on human prostate stem/progenitor cells and prostate cancer risk. Mol. Cell. Endocrinol. 354, 63–73. 10.1016/J.MCE.2011.08.032

28. Huth, H.W., Castro-Gomes, T., de Goes, A.M., Ropert, C., 2021. Translocation of intracellular CD24 constitutes a triggering event for drug resistance in breast cancer. Sci. Rep. 11. 10.1038/S41598-021-96449-7

29. IARC, 2020. Global Cancer Observatory [WWW Document]. URL https://gco.iarc.fr/ (accessed 1.31.22).

30. Ionkina, A.A., Balderrama-Gutierrez, G., Ibanez, K.J., Phan, S.H.D., Cortez, A.N., Mortazavi, A., Prescher, J.A., 2021. Transcriptome analysis of heterogeneity in mouse model of metastatic breast cancer. Breast Cancer Res. 23. 10.1186/S13058-021-01468-X

31. Jang, M.H., Kim, H.J., Kim, E.J., Chung, Y.R., Park, S.Y., 2015. Expression of epithelial– mesenchymal transition-related markers in triple-negative breast cancer: ZEB1 as a potential biomarker for poor clinical outcome. Hum. Pathol. 46, 1267–1274. 10.1016/J.HUMPATH.2015.05.010

32. Jin, F., Wu, Z., Hu, X., Zhang, J., Gao, Z., Han, X., Qin, J., Li, C., Wang, Y., 2019. The PI3K/Akt/GSK-3β/ROS/eIF2B pathway promotes breast cancer growth and metastasis via suppression of NK cell cytotoxicity and tumor cell susceptibility. Cancer Biol. Med. 16, 38–54. 10.20892/J.ISSN.2095-3941.2018.0253

33. Jones, M.L., 2020. Histotechnology a self instructional text, 5th edition. J. Histotechnol. 43, 210–210. 10.1080/01478885.2020.1828682

34. Kolak, A., Kamińska, M., Sygit, K., Budny, A., Surdyka, D., Kukiełka-Budny, B., Burdan, F., 2017. Primary and secondary prevention of breast cancer. Ann. Agric. Environ. Med. 24, 549–553. 10.26444/AAEM/75943

35. Lasagna, M., Hielpos, M.S., Ventura, C., Mardirosian, M.N., Martín, G., Miret, N., Randi, A., Núñez, M., Cocca, C., 2020. Chlorpyrifos subthreshold exposure induces epithelial– mesenchymal transition in breast cancer cells. Ecotoxicol. Environ. Saf. 205. 10.1016/j.ecoenv.2020.111312

36. Lasagna, M., Ventura, C., Hielpos, M.S., Mardirosian, M.N., Martín, G., Miret, N., Randi, A., Núñez, M., Cocca, C., 2022. Endocrine disruptor chlorpyrifos promotes migration, invasion, and stemness phenotype in 3D cultures of breast cancer cells and induces a wide range of pathways involved in cancer progression. Environ. Res. 204. 10.1016/J.ENVRES.2021.111989

37. Lee, H.M., Hwang, K.A., Choi, K.C., 2017. Diverse pathways of epithelial mesenchymal transition related with cancer progression and metastasis and potential effects of endocrine disrupting chemicals on epithelial mesenchymal transition process. Mol. Cell. Endocrinol. 457, 103–113. 10.1016/j.mce.2016.12.026

38. Liu, S., Cong, Y., Wang, D., Sun, Y., Deng, L., Liu, Y., Martin-Trevino, R., Shang, L., McDermott, S.P., Landis, M.D., Hong, S., Adams, A., D’Angelo, R., Ginestier, C., Charafe-Jauffret, E., Clouthier, S.G., Birnbaum, D., Wong, S.T., Zhan, M., Chang, J.C., Wicha, M.S., 2014. Breast cancer stem cells transition between epithelial and mesenchymal states reflective of their normal counterparts. Stem Cell Reports 2, 78–91. 10.1016/J.STEMCR.2013.11.009/ATTACHMENT/7DD43D78-9CFB-42C2-A1AC-465B45012E89/MMC1.PDF

39. Liu, S., Cong, Y., Wang, D., Sun, Y., Deng, L., Liu, Y., Martin-Trevino, R., Shang, L., McDermott, S.P., Landis, M.D., Hong, S., Adams, A., D’Angelo, R., Ginestier, C., Charafe-Jauffret, E., Clouthier, S.G., Birnbaum, D., Wong, S.T., Zhan, M., Chang, J.C., Wicha, M.S., 2013. Breast cancer stem cells transition between epithelial and mesenchymal states reflective of their normal counterparts. Stem cell reports 2, 78–91. 10.1016/J.STEMCR.2013.11.009

40. McLachlan, J., 2016. Environmental signaling: from environmental estrogens to endocrine– disrupting chemicals and beyond. Andrology. 10.1111/andr.12206

41. Palacios-Arreola, M.I., Moreno-Mendoza, N.A., Nava-Castro, K.E., Segovia-Mendoza, M., Perez-Torres, A., Garay-Canales, C.A., Morales-Montor, J., 2022. The Endocrine Disruptor Compound Bisphenol-A (BPA) Regulates the Intra-Tumoral Immune Microenvironment and Increases Lung Metastasis in an Experimental Model of Breast Cancer. Int. J. Mol. Sci. 23. 10.3390/IJMS23052523

42. Palomeras, S., Ruiz-Martínez, S., Puig, T., 2018. Targeting Breast Cancer Stem Cells to Overcome Treatment Resistance. Molecules 23. 10.3390/MOLECULES23092193

43. Park, M.S., Dong, S.M., Kim, B.R., Seo, S.H., Kang, S., Lee, E.J., Lee, S.H., Rho, S.B., 2014. Thioridazine inhibits angiogenesis and tumor growth by targeting the VEGFR– 2/PI3K/mTOR pathway in ovarian cancer xenografts. Oncotarget 5, 4929–4934. 10.18632/ONCOTARGET.2063

44. Peitzsch, C., Tyutyunnykova, A., Pantel, K., Dubrovska, A., 2017. Cancer stem cells: The root of tumor recurrence and metastases. Semin. Cancer Biol. 44, 10–24. 10.1016/J.SEMCANCER.2017.02.011

45. Pontillo, C.A., Rojas, P., Chiappini, F., Sequeira, G., Cocca, C., Crocci, M., Colombo, L., Lanari, C., Kleiman de Pisarev, D., Randi, A., 2013. Action of hexachlorobenzene on tumor growth and metastasis in different experimental models. Toxicol. Appl. Pharmacol. 268, 331–342. 10.1016/j.taap.2013.02.007

46. Ramos Nieto, M.R., Lasagna, M., Cao, G., Álvarez, G., Santamaria, C., Rodriguez Girault, M.E., Bourguignon, N., Di Giorgio, N., Ventura, C., Mardirosian, M., Rodriguez, H., Lux-Llantos, V., Cocca, C., Núñez, M., 2021. Chronic exposure to low concentrations of chlorpyrifos affects normal cyclicity and histology of the uterus in female rats. Food Chem. Toxicol. 156, 112515. 10.1016/J.FCT.2021.112515

47. Sidney, L.E., Branch, M.J., Dunphy, S.E., Dua, H.S., Hopkinson, A., 2014. Concise Review: Evidence for CD34 as a Common Marker for Diverse Progenitors. Stem Cells 32, 1380–1389. 10.1002/STEM.1661

48. Sung, H., Ferlay, J., Siegel, R.L., Laversanne, M., Soerjomataram, I., Jemal, A., Bray, F., 2021. Global cancer statistics 2020: GLOBOCAN estimates of incidence and mortality worldwide for 36 cancers in 185 countries. CA. Cancer J. Clin. 10.3322/caac.21660

49. Takebe, N., Harris, P.J., Warren, R.Q., Ivy, S.P., 2011. Targeting cancer stem cells by inhibiting Wnt, Notch, and Hedgehog pathways. Nat. Rev. Clin. Oncol. 8, 97–106. 10.1038/NRCLINONC.2010.196

50. Tang, D., Chen, M., Huang, X., Zhang, Guicheng, Zeng L., Zhang, Guangsen, Wu, S., Wang, Y., 2023. SRplot: A free online platform for data visualization and graphing. PLoS One 18. 10.1371/JOURNAL.PONE.0294236

51. Thiery, J.P., Acloque, H., Huang, R.Y.J., Nieto, M.A., 2009. Epithelial-Mesenchymal Transitions in Development and Disease. Cell. 10.1016/j.cell.2009.11.007

52. Thomas Zoeller, R., Brown, T.R., Doan, L.L., Gore, A.C., Skakkebaek, N.E., Soto, A.M., Woodruff, T.J., Vom Saal, F.S., 2012. Endocrine-disrupting chemicals and public health protection: A statement of principles from the Endocrine Society. Endocrinology 153, 4097–4110. 10.1210/en.2012-1422

53. Tuomela, J., Härkönen, P., 2014. Tumor models for prostate cancer exemplified by fibroblast growth factor 8-induced tumorigenesis and tumor progression. Reprod. Biol. 14, 16–24. 10.1016/J.REPBIO.2014.01.002

54. U.S. National Academy of Sciences, 2011. GUIDE LABORATORY ANIMALS FOR THE CARE AND USE OF Eighth Edition Committee for the Update of the Guide for the Care and Use of Laboratory Animals Institute for Laboratory Animal Research Division on Earth and Life Studies.

55. Vassalli, G., 2019. Aldehyde Dehydrogenases: Not Just Markers, but Functional Regulators of Stem Cells. Stem Cells Int. 2019. 10.1155/2019/3904645

56. Ventura, C., Nieto, M., Bourguignon, N., Lux-Lantos, V., Rodriguez, H., Cao, G., Randi, A., Cocca, C., Núñez, M., 2016. Pesticide chlorpyrifos acts as an endocrine disruptor in adult rats causing changes in mammary gland and hormonal balance. J. Steroid Biochem. Mol. Biol. 156, 1–9. 10.1016/j.jsbmb.2015.10.010

57. Ventura, C., Núñez, M., Miret, N., Martinel Lamas, D., Randi, A., Venturino, A., Rivera, E., Cocca, C., 2012. Differential mechanisms of action are involved in chlorpyrifos effects in estrogen-dependent or –independent breast cancer cells exposed to low or high concentrations of the pesticide. Toxicol. Lett. 213, 184–193. 10.1016/j.toxlet.2012.06.017

58. Ventura, C., Venturino, A., Miret, N., Randi, A., Rivera, E., Núñez, M., Cocca, C., 2015. Chlorpyrifos inhibits cell proliferation through ERK1/2 phosphorylation in breast cancer cell lines. Chemosphere 120, 343–350. 10.1016/j.chemosphere.2014.07.088

59. Ventura, C., Zappia, C.D., Lasagna, M., Pavicic, W., Richard, S., Bolzan, A.D., Monczor, F., Núñez, M., Cocca, C., 2019. Effects of the pesticide chlorpyrifos on breast cancer disease. Implication of epigenetic mechanisms. J. Steroid Biochem. Mol. Biol. 186, 96–104. 10.1016/j.jsbmb.2018.09.021

60. Wels, C., Joshi, S., Koefinger, P., Bergler, H., Schaider, H., 2011. Transcriptional activation of ZEB1 by Slug leads to cooperative regulation of the epithelial-mesenchymal transition-like phenotype in melanoma. J. Invest. Dermatol. 131, 1877–1885. 10.1038/JID.2011.142

61. Xu, X., Zhang, M., Xu, F., Jiang, S., 2020. Wnt signaling in breast cancer: biological mechanisms, challenges and opportunities. Mol. Cancer 2020 191 19, 1–35. 10.1186/S12943-020-01276-5

62. Yang, X.R., Chang-Claude, J., Goode, E.L., Couch, F.J., Nevanlinna, H., Milne, R.L., Gaudet, M., Schmidt, M.K., Broeks, A., Cox, A., Fasching, P.A., Hein, R., Spurdle, A.B., Blows, F., Driver, K., Flesch-Janys, D., Heinz, J., Sinn, P., Vrieling, A., Heikkinen, T., Aittomäki, K., Heikkilä, P., Blomqvist, C., Lissowska, J., Peplonska, B., Chanock, S., Figueroa, J., Brinton, L., Hall, P., Czene, K., Humphreys, K., Darabi, H., Liu, J., Van ‘T Veer, L.J., Van Leeuwen, F.E., Andrulis, I.L., Glendon, G., Knight, J.A., Mulligan, A.M., O’Malley, F.P., Weerasooriya, N., John, E.M., Beckmann, M.W., Hartmann, A., Weihbrecht, S.B., Wachter, D.L., Jud, S.M., Loehberg, C.R., Baglietto, L., English, D.R., Giles, G.G., McLean, C.A., Severi, G., Lambrechts, D., Vandorpe, T., Weltens, C., Paridaens, R., Smeets, A., Neven, P., Wildiers, H., Wang, X., Olson, J.E., Cafourek, V., Fredericksen, Z., Kosel, M., Vachon, C., Cramp, H.E., Connley, D., Cross, S.S., Balasubramanian, S.P., Reed, M.W.R., Dörk, T., Bremer, M., Meyer, A., Karstens, J.H., Ay, A., Park-Simon, T.W., Hillemanns, P., Arias Pérez, J.I., Rodríguez, P.M., Zamora, P., Benítez, J., Ko, Y.D., Fischer, H.P., Hamann, U., Pesch, B., Brüning, T., Justenhoven, C., Brauch, H., Eccles, D.M., Tapper, W.J., Gerty, S.M., Sawyer, E.J., Tomlinson, I.P., Jones, A., Kerin, M., Miller, N., McInerney, N., Anton– Culver, H., Ziogas, A., Shen, C.Y., Hsiung, C.N., Wu, P.E., Yang, S.L., Yu, J.C., Chen, S.T., Hsu, G.C., Haiman, C.A., Henderson, B.E., Le Marchand, L., Kolonel, L.N., Lindblom, A., Margolin, S., Jakubowska, A., Lubiński, J., Huzarski, T., Byrski, T., Górski, B., Gronwald, J., Hooning, M.J., Hollestelle, A., Van Den Ouweland, A.M.W., Jager, A., Kriege, M., Tilanus-Linthorst, M.M.A., Collée, M., Wang-Gohrke, S., Pylkäs, K., Jukkola-Vuorinen, A., Mononen, K., Grip, M., Hirvikoski, P., Winqvist, R., Mannermaa, A., Kosma, V.M., Kauppinen, J., Kataja, V., Auvinen, P., Soini, Y., Sironen, R., Bojesen, S.E., Dynnes Ørsted, D., Kaur-Knudsen, D., Flyger, H., Nordestgaard, B.G., Holland, H., Chenevix-Trench, G., Manoukian, S., Barile, M., Radice, P., Hankinson, S.E., Hunter, D.J., Tamimi, R., Sangrajrang, S., Brennan, P., McKay, J., Odefrey, F., Gaborieau, V., Devilee, P., Huijts, P.E.A., Tollenaar, R., Seynaeve, C., Dite, G.S., Apicella, C., Hopper, J.L., Hammet, F., Tsimiklis, H., Smith, L.D., Southey, M.C., Humphreys, M.K., Easton, D., Pharoah, P., Sherman, M.E., Garcia-Closas, M., 2011. Associations of breast cancer risk factors with tumor subtypes: a pooled analysis from the Breast Cancer Association Consortium studies. J. Natl. Cancer Inst. 103, 250–263. 10.1093/JNCI/DJQ526

63. Yoshino, I., Kometani, T., Shoji, F., Osoegawa, A., Ohba, T., Kouso, H., Takenaka, T., Yohena, T., Maehara, Y., 2007. Induction of epithelial-mesenchymal transition-related genes by benzo[a]pyrene in lung cancer cells. Cancer 110, 369–374. 10.1002/CNCR.22728

64. Yu, C.C., Chang, Y.C., 2013. Enhancement of cancer stem-like and epithelial– mesenchymal transdifferentiation property in oral epithelial cells with long-term nicotine exposure: reversal by targeting SNAIL. Toxicol. Appl. Pharmacol. 266, 459–469. 10.1016/J.TAAP.2012.11.023

65. Zárate, L. V., Pontillo, C.A., Español, A., Miret, N. V., Chiappini, F., Cocca, C., Álvarez, L., de Pisarev, D.K., Sales, M.E., Randi, A.S., 2020. Angiogenesis signaling in breast cancer models is induced by hexachlorobenzene and chlorpyrifos, pesticide ligands of the aryl hydrocarbon receptor. Toxicol. Appl. Pharmacol. 401. 10.1016/J.TAAP.2020.115093

66. Zhang, Z., Fang, T., Lv, Y., 2022. Prognostic and clinicopathological value of Slug protein expression in breast cancer: a systematic review and meta-analysis. World J. Surg. Oncol. 20. 10.1186/S12957-022-02825-6

67. Zhao, P., Xu, Y., Wei, Y., Qiu, Q., Chew, T.L., Kang, Y., Cheng, C., 2016. The CD44s splice isoform is a central mediator for invadopodia activity. J. Cell Sci. 129, 1355– 1365. 10.1242/JCS.171959

68. Zucchini-Pascal, N., Peyre, L., de Sousa, G., Rahmani, R., 2012. Organochlorine pesticides induce epithelial to mesenchymal transition of human primary cultured hepatocytes. Food Chem. Toxicol. 50, 3963–3970. 10.1016/J.FCT.2012.08.009

## References

1. Hair et al (2020). Res Integ Peer Rev. doi: 10.1186/s41073-019-0069-3

2. Tihanyi et al (2019). J Surg Res. doi: 10.1016/j.jss.2018.10.038

3. Zhao et al (2020). BMC Vet Res. doi: 10.1186/s12917-020-02664-1

4. Han et al (2017). Plos One. doi: 10.1371/journal.pone.0183591

5. Chatzimanouil et al (2019). J Am Soc Nephrol. doi: 10.1681/ASN.2018050515

6. Leung et al (2018). Plos One. doi: 10.1371/journal.pone.0197882

